# CRISPR-enabled control of gene expression sets the isotopic composition of microbial methane

**DOI:** 10.1101/2024.10.28.620722

**Authors:** Jonathan Gropp, Markus Bill, Max K. Lloyd, Rebekah Stein, Dipti D. Nayak, Daniel A. Stolper

**Affiliations:** Department of Earth and Planetary Science, University of California; Berkeley, Berkeley, CA, USA; Department of Molecular and Cell Biology, University of California; Berkeley, Berkeley, CA, USA; Department of Earth and Environmental Sciences, LBNL; CA, USA; Department of Geosciences, Pennsylvania State University; University Park, PA, USA

## Abstract

The stable isotopic composition of biogenic methane varies substantially in the environment and is routinely used to fingerprint its source. However, the underlying cause of this variation is debated. Here, we experimentally manipulate the growth rate of the model methanogen, *Methanosarcina acetivorans*, using CRISPR mutagenesis to generate a tunable version of the key and final enzyme in methanogenesis, methyl-coenzyme M reductase (MCR). We demonstrate that the carbon and hydrogen isotopic composition of methane change as a function of MCR expression and growth rate. Using an isotope enabled metabolic model we show that these changes stem from a substrate-independent increase in reversibility of methanogenic enzymes. Overall, these data provide a novel framework for calibrating growth coupled changes in the isotopic composition of biogenic methane.

## Main text

Methane is a potent greenhouse gas and energy resource that is generated mostly via microbial methanogenesis and catagenesis of organic matter. The stable carbon and hydrogen isotopic compositions (^13^C/^12^C and D/H ratios, denoted as *δ*^13^C and *δ*D, respectively, where D≡^2^H and H≡^1^H) of methane sources vary significantly and, as such, are commonly used to fingerprint gas origins (*1*–*3*). We focus here on the isotopic composition of biogenic methane, which varies significantly in nature (Fig. 1A). For the past half century, this variance has been assumed to be set the metabolic pathway of methanogenesis (e.g., from H_2_+CO_2_, acetate, or methylated compounds) due to irreversible, pathway-specific kinetic isotope effects (*1*) (Fig. 1A) — we term this the “substrate hypothesis”.

**Figure 1:**
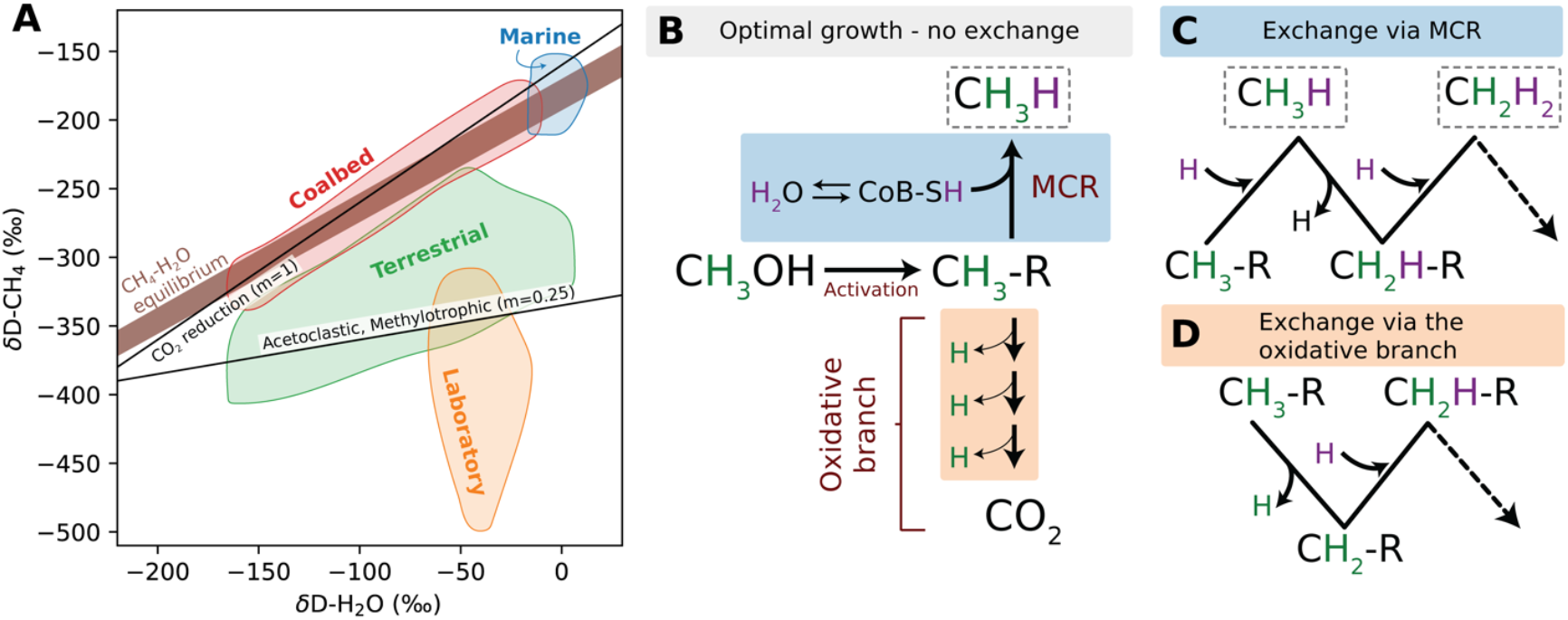
Hydrogen exchange during methanogenesis. (A) The hydrogen isotopic composition of biogenic methane (δD-CH_4_) vs. water (δD-H_2_O) in environmental samples and laboratory cultures. The brown polygon is the CH_4_-H_2_O hydrogen isotopic equilibrium between 0°C (top) and 50°C (bottom) (*6*), the black solid lines represent the canonical end-members for hydrogenotrophic and methylotrophic or acetoclastic (slope of 1 and 0.25, respectively, Ref. (*1*)). Data bounds based on compilation in Ref. (*6*). (B) A schematic view of canonical irreversible methylotrophic methanogenesis. A methyl group from the substrate is activated to an intermediate in the methanogenic pathway (CH_3_-R), which is then either reduced to methane by methyl-coenzyme M reductase (MCR) or oxidized to CO_2_ via the oxidative branch. The hydrogen atoms are colored by their source (methyl-derived H is green, water derived H is purple). (C-D) Potential hydrogen exchange reactions during CH_3_-S-CoM reduction to methane by MCR (C) or during CH_3_-S-CoM oxidation via the oxidative branch (D). Reversible reactions, presented by the zig-zagging arrows, replace methyl-derived hydrogen in methane with water derived hydrogen.

However, substrate availability, i.e., the concentration of reactants used for methanogenesis, has long been known to correlate with the isotopic composition of microbial methane (*4, 5*). Specifically, microbial methane derived from environments with low substrate availability (e.g., deep-sea sediments) appears in isotopic equilibrium with in-situ CO_2_ for carbon isotopes and H_2_O for hydrogen isotopes, and within doubly-substituted “clumped” isotopologues (*6, 7*). In contrast, environments that have abundant substrate concentrations (e.g., cow rumens and lakes) yield microbial methane out of isotopic equilibrium, with the highest disequilibrium seen in substrate-rich laboratory cultures (Fig. 1A) (e.g., *4, 8, 9*). One explanation for the relationship between substrate availability and isotopic compositions is that the enzymes of methanogenesis are reversible and the degree of reversibility (i.e., the relative reverse/forward flux of the enzyme) is a function of free energy (*Δ*G) available to drive methanogenesis (*4, 5, 10*–*12*). Per this model, termed here the “reversibility hypothesis”, at low substrate availability, the *Δ*G of methanogenesis is closer to equilibrium, promoting high reversibility and thus isotopic exchange between hydrogen from water and carbon from CO_2_ with methane. When substrate is abundant, methanogenesis is more energetically favorable (*Δ*G <<<0), and enzymes act irreversibly and express kinetic isotope effects (*4, 5*).

If correct, the reversibility hypothesis requires a fundamental change in the use of stable isotopes to fingerprint environmental methane sources. However, experimental evidence for this hypothesis is limited, in part because laboratory manipulations of methanogens typically occur under high substrate concentrations to accelerate growth. The few prior experiments that attempted to vary the *Δ*G of methanogenesis led to partial C isotope exchange between methane and CO_2_ and no detectable H isotope exchange between methane and water, or among the clumped methane isotopologues (*4, 5, 13, 14*). These observations are consistent with trace methane oxidation (<0.1% of methanogenesis flux) during methanogenic growth under laboratory conditions (*15*–*17*). Modeling studies have also predicted that H isotope exchange in methanogenesis from H_2_+CO_2_ would only be observed in low, nano-molar H_2_ levels, limiting the feasibility of such measurement in lab cultures (*12, 14*). As a result, some researchers have proposed that the near-equilibrium isotopic signatures of microbial methane in substrate-limited environments are instead potentially due to anaerobic methanotrophy, which known to catalyze isotope exchange in both hydrogen and carbon isotopes between methane, water, and CO_2_ (*17*–*20*).

### Isotopic approach for determining enzymatic reversibility of methanogenesis

Here we explore the reversibility hypothesis using a new and distinct approach: we do not attempt to set the *Δ*G of methanogenesis by changing the substrate concentration but, instead, we control the expression of the key and final enzyme of methanogenesis — methyl-coenzyme M reductase (MCR). Inducible expression of MCR is possible due to genomic manipulations facilitated by the development of CRISPR-Cas9 genome editing tools in the model methanogen *Methanosarcina acetivorans* (*21*). This technique allows us to directly test whether changes in enzyme expression, and consequent changes in growth rate, impact the reversibility of methanogenesis as measured by the isotopic composition of carbon and hydrogen in methane. We focus on MCR as it catalyzes the last step of methane production in methanogenic archaea and is shared among all methanogenic pathways (*22*). Importantly, our approach has a unique advantage as it sidesteps uncontrolled and unknown changes in the metabolism of the organism that occur at different substrate concentrations in the laboratory or environment. Instead, through the application of inducible gene expression to MCR, we can directly control methanogenic growth (*23*).

We control expression of the chromosomal MCR operon (*mcr*) via the established tetracycline-inducible gene expression system (*24*) developed specifically for methanogens by (*23*). This system allows for controlled changes in *mcr* expression levels relative to the wild-type strain and has been shown to limit methanogenic growth when <10 μg/mL of tetracycline is present in the growth medium. We experimentally test for changes in reversibility of methanogenesis as a function of *mcr* expression using the hydrogen isotopic composition of methane (δD-CH_4_, where 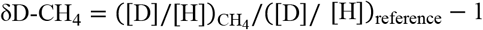 in permil (‰) units). We favor hydrogen isotopes because there are multiple enzymatically catalyzed steps in methanogenesis that, if operated in reverse, will lead to hydrogen isotope exchange with water (Fig. 1B-D). In contrast, the carbon isotopic exchange (δ^13^C-CH_4_, where 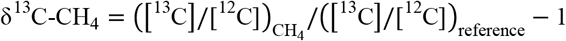 in ‰) between methane and CO_2_ requires every enzymatic step in methanogenesis to operate reversibly. We conducted experiments on methylated substrates (methanol and acetate) as they provide a straightforward way to probe reversibility: under fully irreversible conditions, three hydrogens of methane derive from the methyl group in the substrate, and one from water (*25, 26*) (Fig. 1B). Conversely, as reversibility in the pathway increases, more water-derived H will be incorporated into the final methane, resulting in a change in δD-CH_4_ relative to that formed in the wild-type. (Fig. 1C-D). This change is quantified by varying the isotopic contrast between the starting substrate methyl group vs. the growth-medium water. The larger the initial contrast, the greater the change in the hydrogen isotopic composition of methane for a given amount of enzymatic reversibility. We create differences in this contrast by varying the δD-H_2_O from −90‰ to +2100‰.

### Lowering mcr expression increases enzymatic reversibility

As expected, growth rate slows down with decreasing tetracycline concentrations, confirming prior work showing *mcr* expression and thus MCR concentrations set growth rates (*23*) (Table S1). Important to the work, the decreases in *mcr* expression and growth rate correlate with changes in resultant δD-CH_4_ (Tables S2-3). Specifically, we observe that decreasing *mcr* expression (and thus growth rate) increases the slope and decreases the y-intercept of δD-CH_4_ as a function of growth medium δD-H_2_O (Fig. 2A,D). This demonstrates for the first time that growth rate influences both the hydrogen and the carbon isotopic compositions of microbial methane. This observation could be explained either by changes in the reversibility of enzymes that alter the extent of exchange between methyl derived C-H bonds and water derived H, or by changes in the expressed (unidirectional) kinetic isotope effects of the enzymes that correlates with *mcr* expression levels. We rule out the latter explanation as the specific enzymes used for methanogenesis do not change with changes in *mcr* expression (*23*) and their intrinsic isotope effects should be conserved. As such we propose and proceed with the interpretation that decreasing *mcr* expression changes reversibility of the methanogenic pathway.

**Figure 2:**
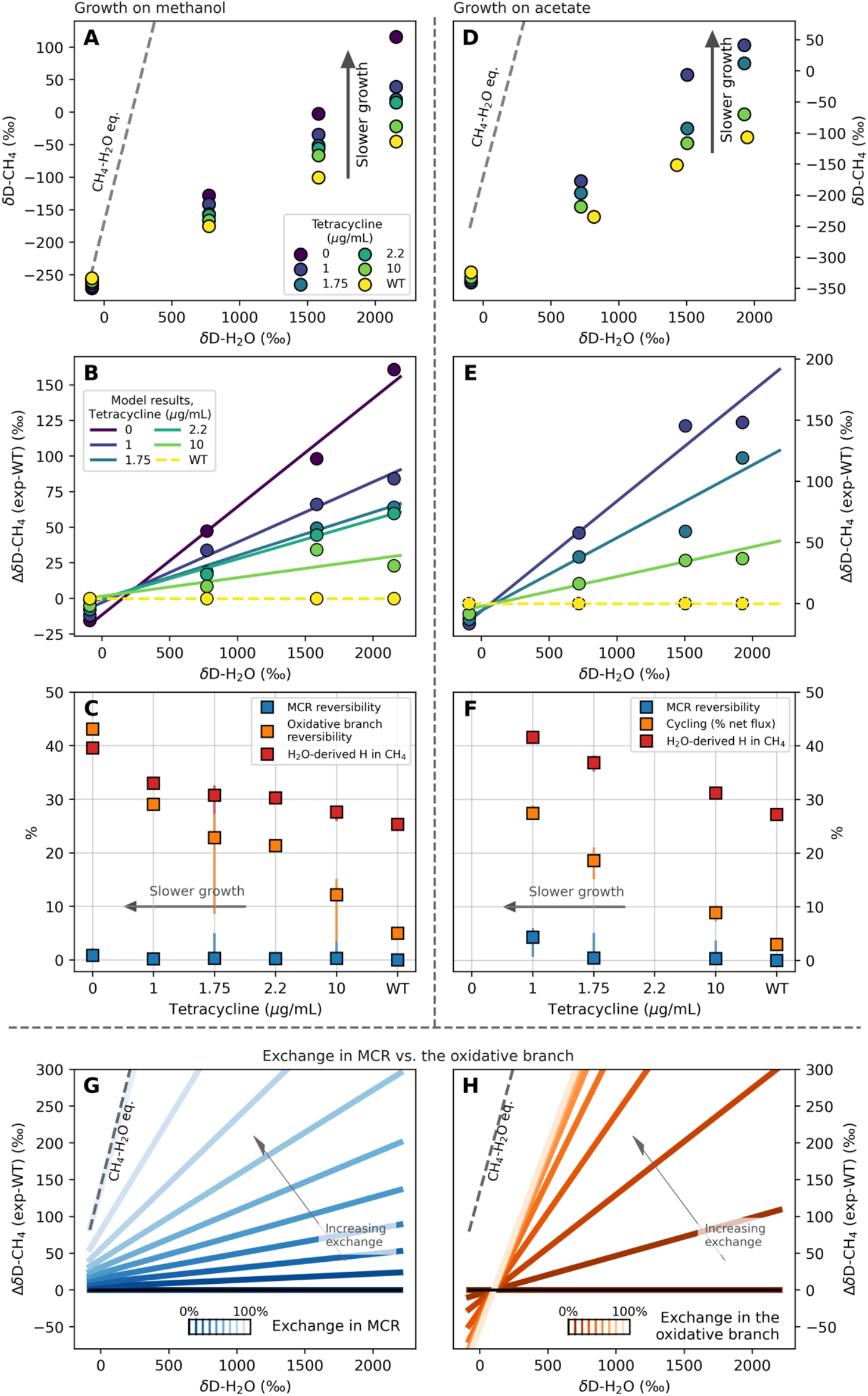
The hydrogen isotopic composition of methane depends on *mcr* expression. The hydrogen isotopic composition of the methane (δD-CH_4_) and source water (δD-H_2_O) for different tetracycline concentrations that represent different concentrations of MCR *in vivo* during growth on (A) methanol and (D) acetate. The gray dashed line denotes CH_4_-H_2_O hydrogen isotopic equilibrium at the growth temperature of 37 °C (*4*). (B, E) Here, the experimental data is presented relative to the best-fit line of the δD-CH_4_ values of the wild-type growth and shown with the fitted modelled results for each *mcr* induction level (where ΔδD-CH_4_ = δD-CH_4(experiment)_–δD-CH_4(WT)_) (solid-colored lines). (C, F) The predicted reversibility degree of the MCR-catalyzed reaction (blue squares) and the oxidative branch (orange squares), and the percentage water-derived hydrogen atoms in methane (red squares). If no H exchange occurs, 25% of the H is expected to come from H_2_O. The error bars represent in panels E and G represent the range that includes 95% of the model results. (G, H) Modeling the δD-CH_4_ as a function of δD-H_2_O for reversibility only in the MCR-catalyzed reaction as normalized to the wild-type condition (G) or only in the oxidative branch (H) in the acetoclastic pathway. The gray dashed line marks the CH_4_-H_2_O hydrogen isotopic equilibrium at 37 °C (*4*), and the dark-gray dotted line is the maximal modelled reversibility obtained in our experiments. The color gradient represents the gradual shift from an irreversible reaction (black line) to a 100% reversible reaction (white line). Notably, an increase in the reversibility of MCR will allow methane to approach CH_4_–H_2_O isotopic equilibrium, while exchange in the oxidative branch increases the slope but also decreases the intercept, such that even at 100% reversibility CH_4_–H_2_O isotopic equilibrium is not possible.

To interpret our results quantitatively, we developed an isotope-enabled model of methanogenesis on methanol and acetate (see Supplementary Material) that includes key reactions in the pathways (Fig. 1B for methanol and Fig. S1B for acetate) and their associated isotope effects. The models include the three key steps of methanogenesis: (*i*) activation of the substrate methyl group to a methylated precursor (CH_3_-R); (*ii*) precursor reduction to methane by MCR; and (*iii*) precursor oxidation via the oxidative branch of methanogenesis to generate electrons for methane production (in methylotrophic methanogenesis) or metabolites for anabolic reactions (in acetoclastic methanogenesis). In order to use the model, we first calibrated the kinetic isotope effects of the enzymes in the pathway based on the wild-type data. Here, we find low degrees of hydrogen isotopic exchange between methyl groups and water due to partial reversibility of the oxidative branch (≈5%, see Supplementary Material). Next, we fit the δD-CH_4_ vs. δD-H_2_O data for each *mcr* expression treatment by assuming that the forward and reverse enzymatic kinetic isotope effects are constant across substrates and treatments and that the only variable parameter is the reversibility of a given enzymatically catalyzed step (Fig. 2B, E). We present here the normalized δD-CH_4_ values of each treatment by subtracting the corresponding δD-CH_4_ values of the WT and denote it as ΔδD-CH_4_ (exp-WT).

For growth on methanol, our model finds that decreased *mcr* expression increases the reversibility of the oxidative branch from ≈5% at wild-type levels up to ≈42% at the lowest expression level (Fig. 2C). In contrast, changes in MCR reversibility were not statistically significant (maximum reversibility of ≈1% at the lowest *mcr* expression) (Fig. 2C). These changes can also be quantified as the amount of water-derived H in the methane, increasing from 25% (effectively irreversible) in the wild-type strain to ≈39% in strains with the highest *mcr* suppression (Fig. 2C).

The underlying basis for why our model only finds an increase in reversibility of the oxidative branch can be visualized based on the δD-CH_4_ vs. δD-H_2_O trends for these two pathways. Specifically, based on the range of initial methanol and water isotopic compositions used, the model suggests that increasing MCR reversibility would rotate and pivot the δD-CH_4_ vs. δD-H_2_O slope towards isotopic equilibrium (such that the intercept increases) with 100% reversibility yield at hydrogen isotopic equilibrium. In contrast, changes in the oxidative branch would lead to a rotation and a change in intercept to lower values (Fig. 2G, H) such that 100% reversibility does not yield equilibrium (as the final H always must come from the action of MCR). Our data are visually consistent with the trajectory predicted by increases in oxidative branch reversibility given the increasing slopes and decreasing intercepts (Fig. 2H) and thus consistent with the general result of the model that most of the changes in reversibility occur in oxidative branch — why this occurs is discussed further below.

Importantly, the model makes an explicit and testable prediction for carbon isotopic of methane during growth on methanol. Increased reversibility in either MCR or the oxidative branch will cause carbon isotope exchange between the precursor methyl groups and the products (CO_2_ and CH_4_). Based on equilibrium isotope effects for different metabolites, increased reversibility in MCR causes δ^13^C-CH_4_ to increase while increased reversibility for the oxidative branch causes δ^13^C-CH_4_ to decrease (see Supplementary Material for explanation). We observe that δ^13^C-CH_4_ decreases with decreasing *mcr* expression by 5.5‰ (Fig. S2). As such, the δ^13^C data corroborate the δD data and our model that decreasing *mcr* expression increases the reversibility of the oxidative branch, but not MCR.

We now turn to experiments with growth on acetate. Here, we build the metabolic model to allow for exchange between methyl groups and H_2_O via the enzymes of the oxidative branch without production of additional CO_2_ (which does not occur based on redox balance) during acetoclastic growth (Fig. S1B). We term this process ‘cycling’ rather than reversibility to highlight that the oxidative branch enzymes must be nearly 100% reversible to maintain redox balance. Although the oxidative branch is commonly not depicted in the metabolic pathway for acetoclastic methanogenesis, it is expressed at low levels in order to generate intermediates for biomass (*27*). Fitting of the model to the experimental data finds that decreasing *mcr* expression causes cycling to increase from ≈3% of the net rate of methanogenesis at wild-type growth to up to ≈30% at the lowest *mcr* expression levels. MCR reversibility also increases, but less so than the oxidative branch, going from 0% at most expression levels up to ≈5% at the lowest *mcr* expression level (Fig. 2F). Finally, the model predicts that there will be no change in the δ^13^C of methane as a function of *mcr* expression as acetoclastic growth does not allow for exchange between the methyl group in acetate and the CO_2_ pool. Our data are consistent with this prediction: *δ*^13^C-CH_4_ stays effectively constant (<1.5‰ difference) across all treatments (Fig. S2).

### Bioenergetic basis for changes in the isotopic composition of methane

A key finding of this study is that we can controllably increase hydrogen isotope exchange between methane and water by titrating MCR expression to lower growth rates. Specifically, we showed that *mcr* suppression and the associated decline in growth rate increases the reversibility of enzymes involved in the oxidative branch of methanogenesis without substantial changes in the reversibility of MCR. Changes in the reversibility of these enzymes must results, at a basic thermodynamic level, from changes in free energy gradients across the various enzymatically controlled reactions. Such changes are not unexpected as it is known that when *mcr* is downregulated, methanogens also change expression of genes in the oxidative branch to reach a new steady state to maintain a constant flux through the metabolic pathway (Fig. S3). We propose that at this new steady state, the ΔG of individual reactions in methanogenesis change as well and that it is these changes that drive the observed reversibility in our experiments. Put another way, although we did not change the overall total ΔG available to cells in the form of substrate concentration, we likely induced changes in the distribution of the ΔG between the methyl transfer, methyl reduction, and methyl oxidation steps (Fig. 1B-D) by modifying *mcr* gene expression.

Overall, our results might still appear somewhat counterintuitive: why would suppression of MCR levels causes significant changes in oxidative branch reversibility but not MCR reversibility? This can be understood by examining the *Δ*G of individual reactions in the methanogenic pathway. By lowering the amount of MCR present, all else being equal, methanogens slow the conversion of CH_3_-S-CoM to methane, leading to an accumulation of CH_3_-S-CoM and coenzyme B, which would alter the redox state of the cell (Fig. 1B), requiring changes in gene expression levels to reach a new steady state (as is known to occur as discussed above). From an energetic standpoint, the oxidative branch has a relatively low overall standard ΔG relative to MCR (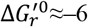versus ≈–25 kJ mol^−1^, respectively) (Table S5). The small ΔG of the oxidative branch is within the range that allows for large changes in reversibility driven by relatively small changes in metabolite concentrations in the oxidative branch (Fig. S4). Additionally, the oxidative branch enzymes are known to function in the reverse direction during methanogenesis on CO_2_+H_2_ (*33*) demonstrating they are readily reversible. In contrast, MCR is effectively irreversible (<0.1%) under typical laboratory growth conditions that lack any extracellular electron acceptors (*16, 28*). As such, our results provide the first experimental evidence that as growth rate changes, methanogens change the reversibility of enzymes involved in the oxidative branch and that this occurs prior to significant changes in MCR reversibility. These changes will modify isotopic composition of methane vs. fully irreversible conditions, without allowing for equilibrium with water or CO_2_ (which requires MCR to reverse).

### Connection to the biogeochemistry of methane

Our work has practical implications for the interpretation of the isotopic composition of environmental methane. Lab-grown pure cultures of methanogens are assumed to generate methane with minimal, if any, isotopic exchange between the methane, water, and the substrates such that the isotopic composition of methane produced is a function of enzyme-controlled kinetic isotopic effects (*1*). This methane has a different isotopic fingerprint relative to biogenic methane from the environment, which is often divided into two groups: (*i*) terrestrial environments, presumed to be dominated by methylotrophic or acetoclastic methanogenesis and (*ii*) marine and coalbed environments, presumed to be dominated by the CO_2_ reductive pathway. An alternative interpretation of these two groups of environmental data is that they instead differ in their degree of isotopic equilibration. Specifically, we propose that the trend towards isotopic equilibrium in terrestrial methane is caused by an elevated degree of isotopic exchange via the oxidative branch as we observed in our experiments (Fig. 3). This incomplete approach to equilibrium would be the result of only partial reversibility of MCR due to partial substrate limitation relative to laboratory conditions. Conversely, in deep-sea sediment methane, extreme substrate limitation suppresses environmental ΔG gradients and slows growth rates to such an extent that MCR becomes significantly reversible, enabling more complete isotopic equilibration. Despite lowering *mcr* expression, we were unable to induce clear increases in the reversibility of MCR — as is required to yield the isotopically equilibrated methane observed in substrate-limited environments. We propose this might be the case because we only slowed growth rates by a factor of ≈5, whereas a much slower growth (by a factor of ≈100) is expected in environments like deep sea sediments where methane is found in isotopic equilibrium (Fig. 3) (*12, 29*). It is likely that such equilibration requires further suppression of MCR activity (and thus growth) beyond what was accomplished in this study and that this suppression would lead to even slower growth rates and drive the δD-CH_4_ towards equilibrium (Fig. 3).

**Figure 3:**
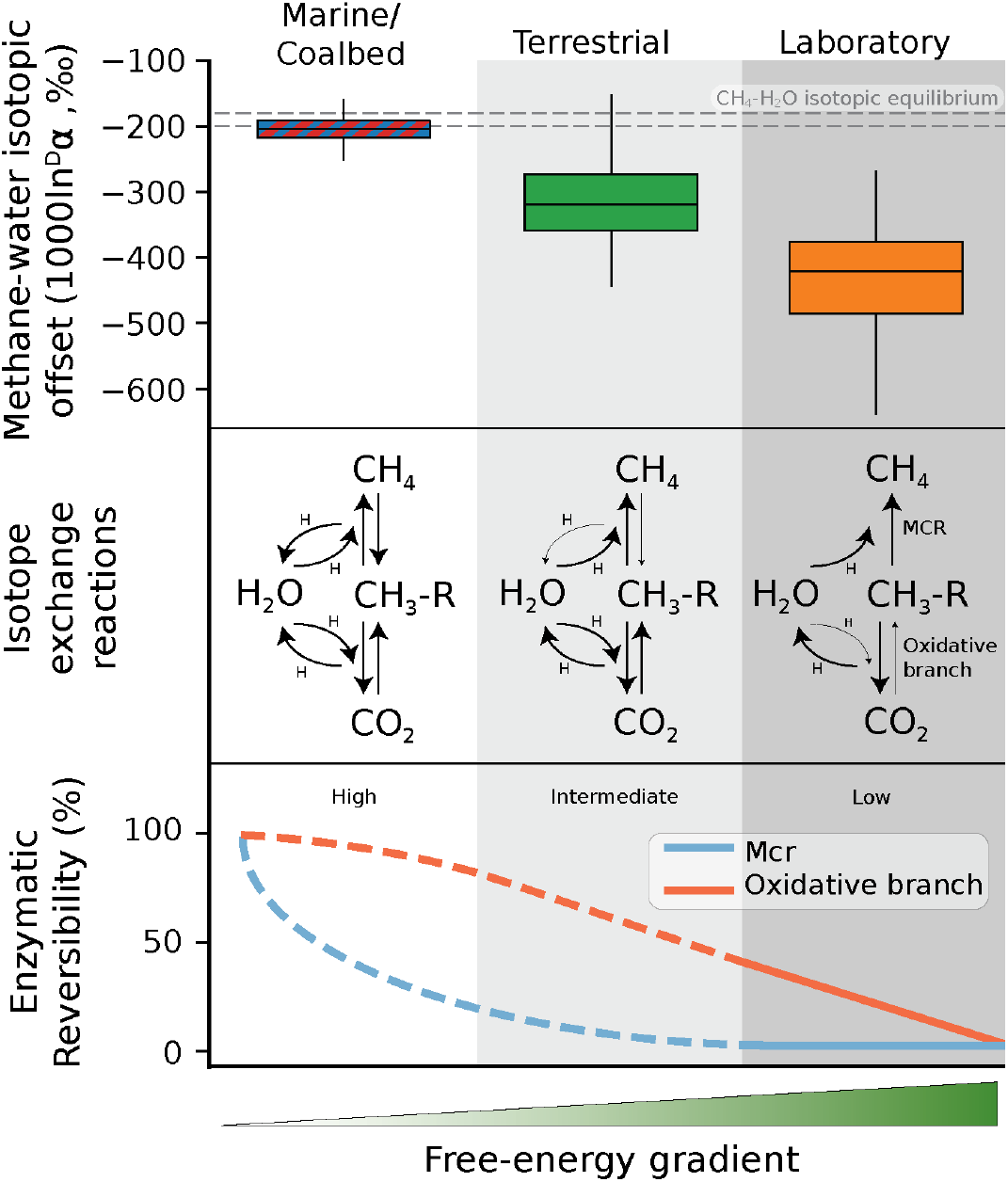
Lower growth rates increase pathway-specific reversibility, ultimately driving CH_4_–H_2_O isotopic composition towards equilibrium. A conceptual illustration of factors that control the isotopic offset between CH_4_ and H_2_O in lab experiments and natural environments. The isotopic offset is presented as 1000 In ^*D*^*α*, where 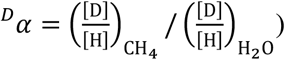. The box-and-whisker plots of compiled data are given in the top pane along with values expected for equilibrium at 50 and 10 °C in the dotted horizontal lines (top and bottom lines, respectively). Our interpretations about the relations between free energy and growth rates to the degree of enzymatically catalyzed isotope-exchange reactions are also given. Light arrows indicate more limited exchange relative to heavier arrows. In our experiments, lower growth rates promoted higher reversibility in the oxidative branch of methylotrophic and acetoclastic methanogenesis and decreased the 1000 In ^*D*^*α* values. Terrestrial environments, in general, have lower 1000 In ^D^*α* values relative to lab experiments, but do not obtain full CH_4_–H_2_O isotopic equilibrium. We propose that this is due to the increasing degree of reversibility in the oxidative branch and a gradual increase in the reversibility of MCR. Environments where growth rates are predicted to be even lower, likely due to low availability of substrates, such as marine sediments and coal bed methane often show full CH_4_–H_2_O isotopic equilibrium, due to high degree of reversibility in all steps of the methanogenic pathway (data in top panel from Ref. (*6*)).

In summary, our work provides the first clear experimental evidence in favor of the “reversibility hypothesis” by showing that the degree of isotopic exchange between methane and water and CO_2_ can be manipulated and controlled. It is not only the metabolic pathways that cause the differences observed in the isotopic composition of environmental methane, but also the physiology of the methanogens as they respond to different environmental constraints (e.g., free energy gradients). Importantly, we do this at the enzymatic level by linking the free energy of specific enzymatic reactions to the overall reversibility of the methanogenic pathway. Taken together, we find strong experimental support for the hypothesis that the isotopic composition of methane is set by variable degree of reversibility regardless of methanogenic substrates involved.

## Acknowledgements

J.G. was supported by the EMBO Postdoctoral Fellowship and by the Institute for Environmental Sustainability (IES). DAS would like to acknowledge support from the Alfred P. Sloan Research Fellowship sponsored by the Sloan Foundation. D.D.N. would like to acknowledge funding from the Searle Scholars Program sponsored by the Kinship Foundation, the Beckman Young Investigator Award sponsored by the Arnold and Mabel Beckman Foundation, the Simons Early Career Investigator in Marine Microbial Ecology and Evolution Award sponsored by the Simons Foundation, the Alfred P. Sloan Research Fellowship sponsored by the Sloan Foundation and the Packard Fellowship in Science and Engineering sponsored by the David and Lucille Packard Foundation. D.D.N is a Chan-Zuckerberg Biohub – San Francisco Investigator.

## Supplementary Materials

### Materials and Methods

#### Strains and Growth Media

Two *Methanosarcina acetivorans* strains were used in this study: (*i*) strain WWM60, wherein the *hpt* locus is replaced with a tetracycline repressor gene (*tetR*) expressed under an *mcrB* promoter from *Methanosarcina barkeri* Fusaro (Δ*hpt*::P*mcrB-tetR*) (*1*). We consider here WWM60 as the “wild type” (WT) *M. acetivorans* strain. (*ii*) Strain DDN032 was generated in the genetic background of WWM60 where the native promoter of the *mcr* operon was replaced by the tetracycline inducible promoter (WWM60-P*mcrB*(*tetO1*)-*mcrBDCGA*) (*2*). Previous work showed a ≈100 fold decrease in *mcr* transcripts, accompanied by a decrease in the methyl coenzyme M reductase (MCR) protein abundance between the highest and lowest concentrations of tetracycline used there: 100 and 1 *μ*g/mL tetracycline. Additionally, across this range, growth rates were reduced by ≈3 folds for growth on methanol and ≈4 folds for growth on acetate (*2*).

Across all treatments and substrates, strains were grown in single-cell morphology at 37 ^°^C without shaking in PIPES-buffered high-salt (HS) liquid medium (*3*) with an over-pressurized He headspace (70 kPa). The medium was supplemented with either 125 mM methanol or 40 mM acetate. Anaerobic, sterile fresh stocks of tetracycline hydrochloride were prepared in deionized water before use and added to final concentrations between 1 to 100 *μ*g/mL.

Each experiment started from a cryo-preserved stock of WWM60 or DDN032 grown on methanol. DDN032 was supplemented with 100 *μ*g/mL tetracycline. Growth was conducted in 26 mL Balch tubes with 10 mL medium and 0–10 *μ*g/mL tetracycline. Prior to inoculation the cells were washed three times PIPES-buffered HS liquid medium to avoid carryover of tetracycline. Each culture used for growth measurement was incubated with cells that would result in an initial optical density at 600 nm (OD_600_) of ≈0.1. We considered an experiment concluded when the cultures were at the their maximal optical density for at least 48 hours. Following the determination that an experiment was concluded, tubes were stored at room temperature. At routine intervals, the genotype of DDN032 was verified by whole-genome sequencing to check for the presence of “escape” mutants which were shown to repress *tetR* and the associated tetracycline-based induction of *mcr* expression, as described previously in Ref. (*2*). No escape mutants were found across all growth experiments that were randomly sampled.

#### Isotopic Analyses

The carbon and hydrogen isotopic compositions (*δ*^13^C and *δ*D, expressed in the VPDB and VSMOW frameworks, respectively) of CH_4_ were analyzed at the Center for Isotope Geochemistry at the Lawrence Berkeley National Laboratory using a gas-chromatograph isotope ratio mass spectrometer (GC-IRMS) system (Thermo Scientific GC TraceGas Ultra system connected to a Thermo Scientific Delta V Plus). The methodology employed here follows that described in (*4*). Briefly, headspace gas was sampled using a gas-tight syringe and injected into a stainless-steel loop (5 *μ*L to 100 5 *μ*L) attached to a 6-port valve (VICI-Valco). The CH_4_ was separated chromatographically on an HP-mole sieve fused silica capillary column (30 m ± 0.32 mm) using helium as carrier gas. Following chromatographic separation, CH_4_ was passed through a ceramic tube at 1020 ^°^C, combusted, and converted to CO_2_ for carbon isotopic measurements, or through a ceramic tube at 1420 ^°^C, pyrolyzed, and converted to H_2_ for hydrogen isotopic measurements. Ceramic tubes were pre-conditioned by injecting 250 *μ*L of pure CH_4_ three times the day before the measurement session. Measured *δ*D and *δ*^13^C values were corrected relative to external natural gas standards acquired from the United States Geological Service HCG-1 (*δ*D-CH_4_ of −64.0 ‰; *δ*^13^C-CH_4_ of −1.51 ‰), HCG-2 (*δ*D-CH_4_ of −183.2 ‰; *δ*^13^C-CH_4_ of −43.09 ‰), and HCG-3 (*δ*D-CH_4_ of −224.3 ‰; *δ*^13^C-CH_4_ of −61.39 ‰) (*5*).

The bulk carbon and hydrogen isotopic composition of the methanol stock was measured following previously described protocols (*6*). Briefly, pure methanol was derivatized to CH_3_I using HI acid, which was then purified using a series of cryogenic and chemical steps. CH_3_I was then converted to chloromethane (CH_3_Cl) using silver chloride (AgCl). Finally, the isotopic composition of CH_3_Cl was measured using a MAT 253 Ultra (Thermo Scientific) at UC Berkeley. The methanol analysis yielded a calibrated *δ*D value of −49.0 ± 0.5 ‰ (2SE) and *δ*^13^C value of −41.67 ± 0.02 ‰ (2SE).

The bulk carbon isotopic composition of acetate was measured at the Center for Isotope Geochemistry at the Lawrence Berkeley National Laboratory. Silver capsules with pure sodium acetate were loaded into a zero blank autosampler connected to an ECS 4010 Elemental Analyzer (EA, Costech Analytical Technologies Inc.) coupled to a Delta V plus IRMS (Thermo Scientific). The carbon isotopic composition of the acetate methyl group was measured using previously described methods (*7*). Briefly, pure sodium acetate was mixed with sodium hydroxide in a 1:10 molar ratio, and pyrolysed at 400 ^°^C under vacuum, converting the methyl carbon to CH_4_ and the carboxyl carbon to CO_2_. The carbon isotopic composition of the methane was measured at the Center for Isotope Geochemistry at the Lawrence Berkeley National Laboratory as described above. The *δ*^13^C value of the methyl group obtained from two individual pyrolysis experiments was −37.1 ± 2.3 ‰ (2SE). The *δ*D value of sodium acetate was determined via high temperature conversion and isotope ratio mass spectrometry at the Pennsylvania State University following standard methods (*8*). Briefly, 250 to 600 *μ*g of the sodium acetate sample were loaded into pressed silver capsules (EA Consumables) and pyrolyzed at 1450 ^°^C in a glassy carbon reactor on a Temperature Conversion Elemental Analyzer device (TC/EA, Thermo Scientific). Evolved H_2_ was entrained in ultra high purity Helium and separated from other gases (CO) on a gas chromatography column packed with molecular sieve held at 70 ^°^C, and introduced into a Thermo Delta XP IRMS instrument through an open split on a Thermo Conflo IV interface. Yields of H_2_ from sodium acetate were between 96% and 103%. *δ*D values were placed on the VSMOW-SLAP scale using a two-point calibration based on four aliquots each of VSMOW (*δ*D of 0 ‰) and VSLAP2 (*δ*D of −427.5 ‰) in sealed silver tubes [USGS Reston, Ref. (*9*)] measured on the same day in the same analytical session. Accuracy was verified by analyzing four aliquots of an additional water standard (UC04) and recovering the accepted value within error (measured *δ*D of 115.1 ± 2.0 ‰ (2SE) vs. accepted value of 114 ‰). Five analyses of the sodium acetate yielded a calibrated *δ*D value of −178.8 ± 0.9 ‰ (2SE).

Waters with varying *δ*D were prepared by gravimetric mixing of 99.999% pure D_2_O with double-distilled H_2_O. Based on this, we generated *δ*D-H_2_O values between ≈ −90 ‰ (Berkeley double distilled tap water) and ≈ +2, 000 ‰. Water hydrogen isotopic compositions were measured at the Center for Stable Isotope Biogeochemistry at UC Berkeley, via conversion of water to H_2_ using a hot chromium reactor unit (Thermo H/Device) connected to a Thermo Delta V Plus isotope ratio mass spectrometer. External water standards of known isotopic composition GFLES-2 (159.9 ‰), GFLES-3 (280.2 ‰), and GFLES-4 (399.8 ‰) (USGS) were used to correct samples to the international VSMOW reference frame.

#### Model

In order to quantitatively interpret the experimental results, we constructed models based on a reaction network of methylotrophic and acetoclastic methanogenesis. These models predict the isotopic composition of methane as a function of the degree of reversibility (*r*_*i*_) in a given pathway (figure S1). Both models include (*i*) a methylated substrate activation step to a methyl precursor (CH_3_-R), (*ii*) reduction of the CH_3_-R to methane, and (*iii*) oxidation of CH_3_-R.

The CH_3_-R reduction and oxidation reactions in our model are assumed to be able to be reversible such that a given reaction can proceed in both forward and reverse directions. Each reaction is assigned with forward and backward gross fluxes (*J*_*i, f*_ and *J*_*i,b*_, respectively), which are related by the degree of reversibility term *r*, where *r*_*i*_ = *J*_*i,b*_/*J*_*i, f*_. This ratio can be related to the thermodynamic drive of the reaction by *r*_*i*_ = exp (ΔG_r_/RT) where ΔG_r_ is the Gibbs free energy of the reaction, R is the ideal gas constant, and T is the temperature (*10*). In this framework, the degree of reversibility for a reaction with a positive net flux (i.e., *J*^+^ > *J*^−^) approaches a value zero when *J*^+^ >>> *J*^−^ (a practically irreversible reaction with a large absolute free energy gradient), or approaches a value of 1 when *J*^−^ ≈ *J*^+^ (a practically fully reversible reaction where the free energy gradient is 0). We limit the reactions here to positive new fluxes, i.e., net methane oxidation is not allowed. The full set of equations are given further below. We first describe various assumptions made in the model before presenting the equations.

For the model, we assume that the methyl substrate activation step (reaction 1 in figure S1) has effectively no reverse flux and is treated as irreversible. This assumption follows predictions from metabolic models that this step is irreversible (*11*). In contrast, all other reactions (Reaction 2-5 in figure S1) are allowed to be reversible with no *a priori* restrictions on their degree of reversibility other than they must be between 0 and 1. We further assume that the degree of reversibility of the reactions in the oxidative branch identical (Reaction #3 to #5 in figure S1). While these reactions probably have different degrees of reversibility (*12*), in this model they are combined together and cannot be isolated. We thus refer here to the “net” reversibility of the oxidative branch, which is calculated following the suggestion for a parallel reaction network where 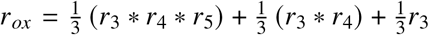. Since we assume *r*_3_ = *r*_4_ = *r*_5_, this can be simplified to 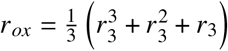 (*13*).

We assume that under irreversible, high absolute free energy growth conditions, the three hydrogen atoms in the substrate methyl group are quantitatively transferred to the methane, and the remaining fourth hydrogen atom is inherited from the water (*14–17*). As such, the maximum ratio of methyl-derived vs. water derived hydrogen in methane is 3:1. We note that there is evidence of partial isotopic exchange between the substrates and a yet unknown intermediate in the MCR catalyzed reaction (*18*). Due to the uncertainty regarding this exchange we do not include it explicitly in the model. We discuss this topic further in the supplementary text. Hydrogen exchange reactions between C-H bonds in precursor molecules or methane with water will alter this 3:1 ratio, increasing the amount of water-derived hydrogen atoms (*17*). It is important to note that while water is the ultimate source of the additional, non-methanol- or acetate-derived hydrogen atoms, two of the exchanged hydrogen atoms are not derived directly from water, but instead via intermediate electron carriers. In the MCR catalyzed reaction a hydrogen atom from coenzyme B (HS-CoB) is added to the three methyl hydrogen atoms of methyl coenzyme M (CH_3_-S-CoM) to form methane. In contrast, in the methylene-tetrahydromethanopterin reductase (MER) and F420-dependent methylene-tetrahydromethanopterin dehydrogenase (MTD) catalyzed reactions (Reaction #3 and #4, respectively, in figure S1) the hydrogen atom is transferred to coenzyme F420. We assume that the hydrogen from these intermediates is derived from water and likely rapidly exchanged and equilibrated with the water.

In the methylotrophic pathway on methanol, CH_3_-S-CoM is disproportionated between the oxidative pathway to CO_2_ and the reductive pathway to methane in a 1:3 ratio (*19*). In the oxidative branch, CH_3_-S-CoM is ultimately fully oxidized to CO_2_ in six consecutive enzymatically-catalyzed reactions. Here, we track the three oxidation reactions where a hydrogen is exchanged with coenzyme F420 (Reaction #3 and #4 in figure S1) or with H_2_O (Reaction #5 in figure S1). We exclude reactions which have only a secondary isotopic effect, where the C-H bond incolved in the reaction is not broken. In the acetoclastic pathway, the methyl precursor methyl-tetrahydromethanopterin (CH_3_-H_4_MPT) is only taken up by the oxidative branch for anabolic reactions, such as for nucleic acid biosynthesis (*20*). Given that nucleic acids are at most ≈10 % of the dry cell weight, and that the total carbon flux to biomass is ≈10 %, the overall anabolic flux through the CH_2_-R and CHR intermediates is only ≈1 % (*21, 22*). Regardless, this is important for our purposes as it means that the oxidative branch, though often not depicted as part of acetoclastic methanogenesis, is operating and thus can allow for hydrogen exchange between C-H bonds in precursor molecules to methane. Based on this, we include for the acetoclastic model a reaction between the CH_3_-R and CH_2_-R (Reaction #3 in figure S1B). We assume that Reaction #3 has a zero-net rate, i.e., the gross forward and backward rates are equal preventing CO_2_ generation occurring (and thus a redox imbalance). We thus define the ‘cycling’ parameter which defines the magnitude of those forward and reverse gross fluxes relative to the net methanogenic flux.

#### Isotopic Mass Balance

We calculate the evolution of the isotopic composition of the substrates and products of the methylotrophic and acetoclastic methanogenesis pathways by forward integrating a set of ordinary differential equations (ODEs, see equations below). These ODEs include time-derivatives of the concentrations and isotopic compositions of the reactant methanol or acetate, and products CO_2_ and CH_4_. We assume a steady-state of all the intermediates is obtained between integration time points (i.e., all the reactions have the same net flux). This allows us to cancel out several parameters in the model such as the concentrations of the intermediates and is the common assumption in models of this sort (*23*). For simplicity, we include only the intermediates that participate in potential isotopic exchange (figure S1). We also include a biomass sink, with an isotopic effect of 1 (i.e., no isotopic effect). We set the biomass sink to 15% of the total flux, which is in the intermediate range for methylotrophic and acetoclastic growth in *Methanosarcina* (*24*). The magnitude of the biomass sink affects the intercept of the *δ*D-CH_4_ vs. *δ*D-H_2_O and not the slope (figure S5). We note that the choice of the exact value of the biomass sink thus affects the absolute value of the fitted isotopic effects (discussed below), but does not change the outcomes of the model.

The gross fluxes in the model are assigned with kinetic isotopic effects (KIEs, 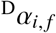 and 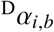, for forward and reverse KIEs, respectively, where 2 denotes deuterium). Here *α* = *k* _*D*_/*k*_*H*_, where *k* _*D*_ and *k*_*H*_ are the normalized rate constants for the reactions with the deuterated and the non-deuterated isotopologues, respectively. The forward and reverse KIEs are related via the temperature-dependent equilibrium isotopic effect 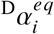, where 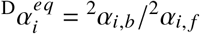. We use the theoretically calculated 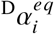 values for the methanogenic pathway to relate the forward and backward KIEs (*25*), thus removing a degree of freedom from the model.

As noted above, the hydrogen atoms in methane are derived either from the substrate or from the medium water (via the electron carries). We assume here that the water hydrogen pool is infinite, and the intracellular and extracellular pools have identical isotopic composition. In other words, the *δ*D-H_2_O values do not change over the course of an experiment. In-vitro studies using purified MCR show effectively immediate isotopic equilibration between H_2_O and the -SH group of HS-CoB, which would lead to H_2_O–HS-CoB hydrogen isotopic equilibrium (*18*). We thus calculate the hydrogen isotopic composition of HS-CoB using the theoretically-calculated isotopic equilibrium fractionation factor between H_2_O and HS-CoB [0.452 at 37 ^°^C, Ref. (*25*)]. Based on theoretical predictions of high reversibility of the reactions that cycle F420 in methylotrophic methanogenesis (*11*), we follow a similar assumption for the isotopic composition of F420 using an equilibrium isotopic fractionation factor of 0.886 between H_2_O and F420 at 37 ^°^C (*25*). Finally, the FMD catalyzed reaction exchanges hydrogen atoms directly with the intracellular water and thus Reaction #5 in figure S1 does not proceed through H-bearing intermediates (*26*).

#### Algorithm for Fitting Model Parameters

The model described above has two groups of free parameters – the hydrogen isotope KIEs and the reversibility of each step. We assume that the KIEs are shared between all experimental conditions (i.e., the enzymes in the organsims share the same KIEs). We allow that the reversibility of each step can differ. Based on this, the differences in the *δ*D-CH_4_ vs. *δ*D-H_2_O are governed by the degree of reversibility of the reactions (and inital *δ*D of the methyl group in methanol and acetate vs. the *δ*D-H_2_O). As such, we can use the model to constrain the reversibility of the various reactions that yield the best fit for the data. Our algorithm and workflow is as follows:

1. The hydrogen isotope KIE for MCR associated with the final H addition to a methyl group to form methane has been measured experimentally (*18*). To our knowledge, no other KIEs have been measured. As such, we follow the approach of Refs. (*12*) and (*27*) for hydrogenotrophic methanogenesis and estimate the KIEs of each reaction (and associated uncertainties). We do this by fitting the *δ*D-CH_4_ vs. *δ*D-H_2_O results of the wild-type *M. acetivorans* WWM60 strain (WT) grown on acetate and methanol (tables S2-S3). We make the following assumptions in determining the KIEs. We assumed that for WT growth, substrate activation and the MCR-catalyzed steps are irreversible (i.e., *r*_1_ = 0 and *r*_2_ = 0). Substrate activation is always assumed to be irreversible (see discussion above). That MCR is irreversible under typical growth conditions is based on the following: (*i*) measurements of MCR reversibility in-vitro shows a backward to forward flux ratio of ≈1/10,000 (*28*); (*ii*) in-vivo isoptope labeling experiments which have detected only trace methane oxidation during methylotrophic growth [< 0.1 %, Ref. (*29*)]; and (*iii*) and that both the activation step and MCR have large-negative standard Gibbs free energies (table S5). We allowed for the possibility of reversibility in the oxidative branch and found that best-fit occurs when this is 5% reversible for WT growth on methanol, and 3% cycling for WT growth on acetate. The prediction of partial reversibility in the oxidative branch at typical methylotrophic and acetoclastic laboratory growth conditions (i.e., optimal conditions) is supported both experimentally (*17, 30*), and theoretically by metabolic models (*11*).
2. To fit the WT growth experiments, we ran the isotopic model 10^5^ times while drawing the KIEs of MCR and the oxidative branch from random uniform distributions with 0.3 ≤ *α* ≤ 1 for primary KIE, where a C-H bond is formed or broken, and 0.7 ≤ *α* ≤ 1 for secondary KIEs where the C-H bond is not affected directly. These ranges represent typical hydrogen isotopic effects for enzymatically catalyzed reactions (*31*). We use a KIE of 0.95 for the methanol activation step. Since the substrates are quantitatively converted to CH_3_-S-CoM without exchange, the magnitude of this KIE does not affect the *δ*D-CH_4_ vs. *δ*D-H_2_O relation. We calculated the sum of squared errors (SSE) of the modeled vs. observed *δ*D-CH_4_ values. The SSE was used as weight for each individual randomly drawn prior KIE to generate the posterior distributions of the KIE values which were then used to fit the experimental data (see the following discussion on the posterior KIE values).
3. To fit the experimental data in different *mcr* expression levels, we then ran the model again using the median posterior KIE values which were now not allowed to change. We then draw the degree of the reversibilities of the MCR catalyzed reaction (*r*_2_) and the oxidative branch (*r*_3_) from a uniform distribution (0 ≤ *r* ≤ 1), to calculate the prior *δ*D-CH_4_ values as a function of *δ*D-H_2_O. We then calculated the SSE separately for each experimental data set (i.e., each tetracycline treatment) relative to the experimentally measured *δ*D-CH_4_ vs. *δ*D-H_2_O values. We found that the ideal fits were obtained when we used the SSE of two data points along the linear regression of the actual data, at a *δ*D-H_2_O of −300 ‰ and +1, 100 ‰. The SSE was used as weight to generate the posterior *r*_2_ and *r*_3_ values (figure 2E,G). The posterior *δ*D-CH_4_ values were calculated by running the model again with the posterior reversibilities and KIEs (figure 2C-D).

#### Model Equations for Methylotrophic Methanogenesis on Methanol

The time derivative of the concentrations and isotopic composition of the metabolites that are in included in the model are as follows:

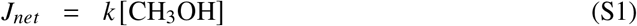

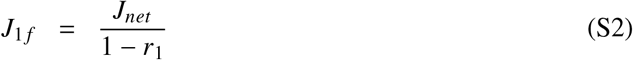

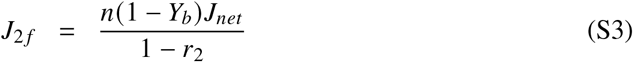

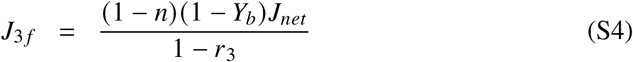

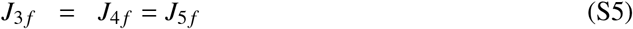

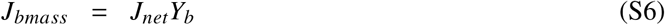

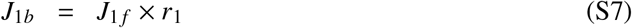

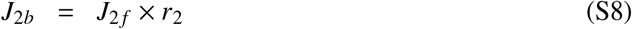

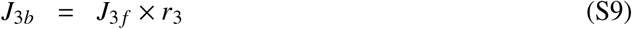

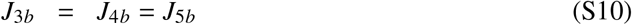

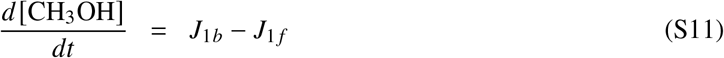

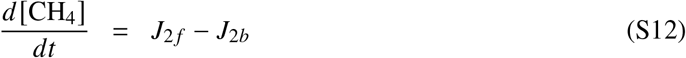

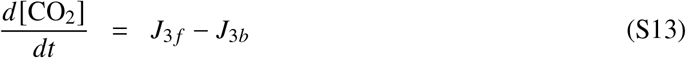

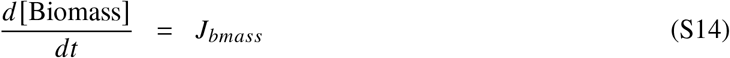

where *J*_*i, f*_ and *J*_*i,b*_ are the forward and backward fluxes of the *i*th reaction. See figure S1 for the reactions and their corresponding indices. *J*_*net*_ is the net rate of methanol consumption, *Y*_*b*_ is the biomass sink, *n* is the fraction of CH_3_R that is reduced to CH_4_, and *r*_*i*_ is the degree of reversibility of the *i*th reaction. The value of *Y*_*b*_ is 0.15, and *n* is 0.75.

We now define the time derivatives of the isotopic ratios of the metabolites in the pathway (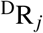 for metabolite *j*). Here, 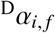 and 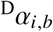 are the forward and backward kinetic isotopic effects (see section for definitions), and the *s* and *p* superscripts denote the secondary and primary hydrogen KIEs.

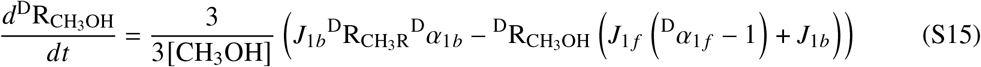

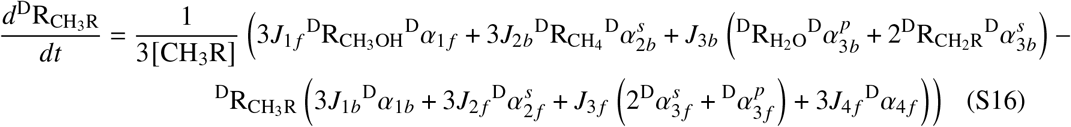

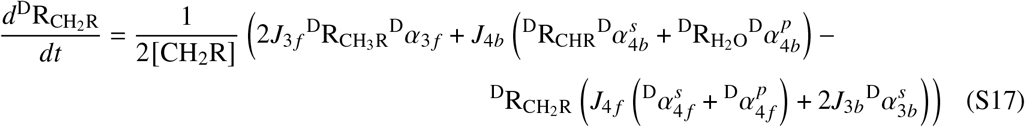

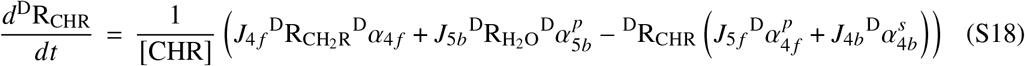

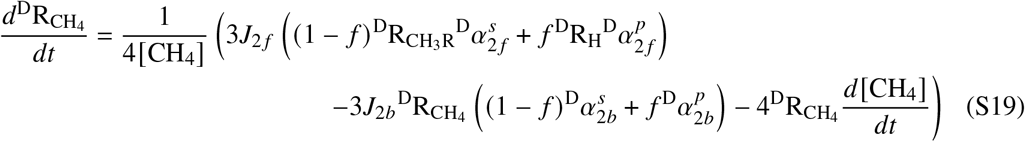

By assuming that in each time point the intermediates obtain a steady state, we can write eq. S16 as:

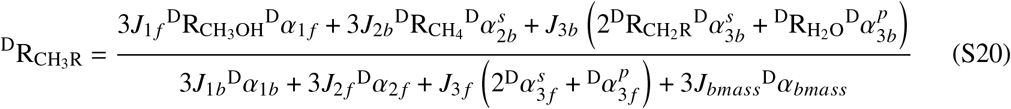

We now express 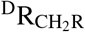 and 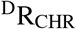 in terms of 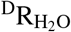 to isolate 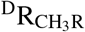, under the assumptions that *J*_3 *f*_ = *J*_4 *f*_ = *J*_5 *f*_, and *r*_3_ = *r*_4_ = *r*_5_. We then use the following identities to isolate the steady-state isotopic ratio of CH_3_R:

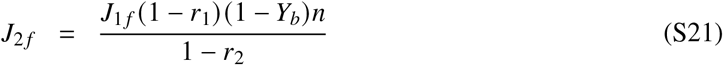

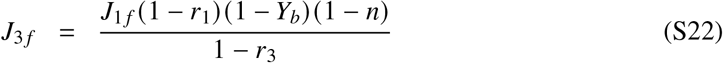

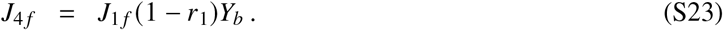

which yields:

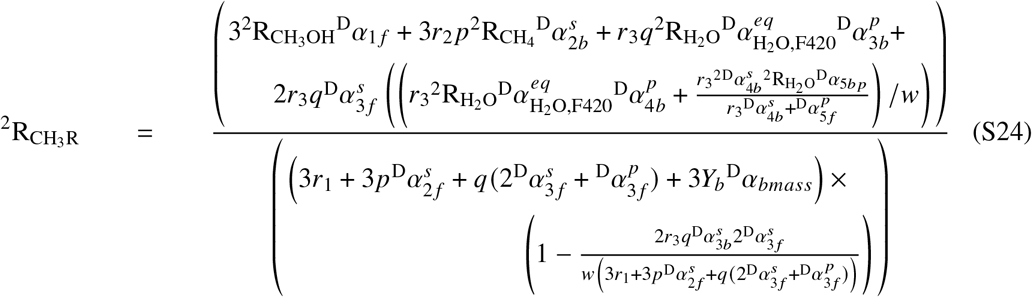

where:

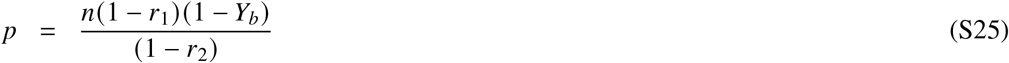

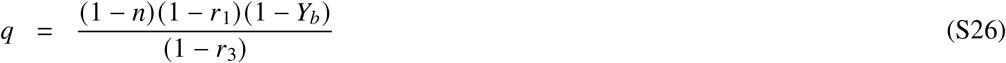

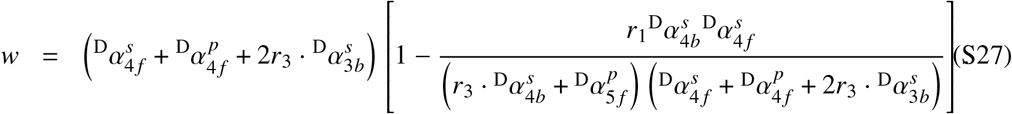

#### Model Equations for Acetoclastic Methanogenesis

We follow a similar derivation for the acetoclastic pathway.

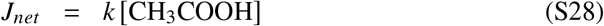

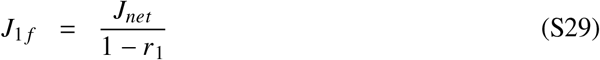

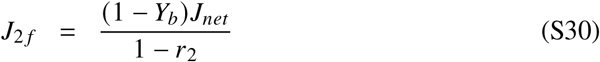

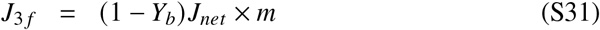

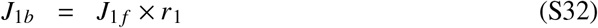

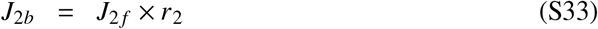

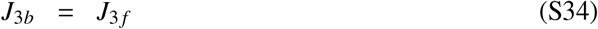

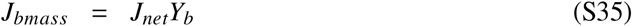

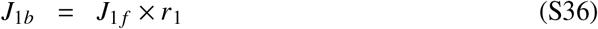

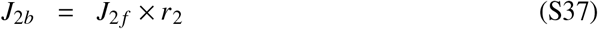

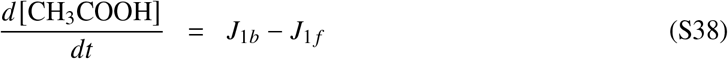

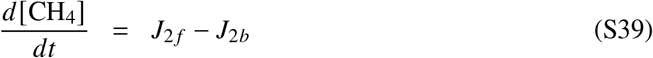

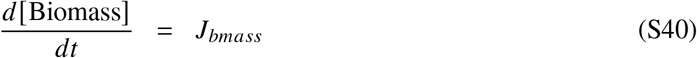

Note that here the third reaction has a zero net flux (i.e., *J*_3 *f*_ = *J*_3*b*_), and is by definition fully reversible. The term *m* refers to the degree of cycling via the relation *J*_3 *f*_ = (1 − *Y*_*b*_) *J*_*net*_ × *m*. Following a similar assumption of steady state between model integration steps we obtain the steady-state isotopic composition of CH_3_R:

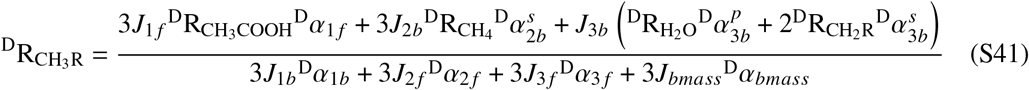

Following the assumption that the oxidative branch in the acetoclastic pathway does have a net flux and is thus at isotopic equilibrium, we set the isotopic composition of 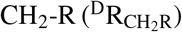 at hydrogen isotopic equilibrium with the water.

#### Calculation of the Fraction of Water-Derived Hydrogen Atoms in CH_4_

We write a set of ordinary differential equations for the hydrogen pools in the model, irrespective of the isotopes:

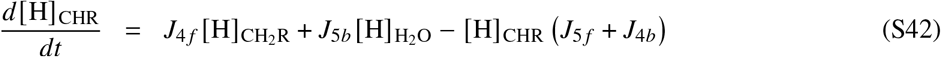

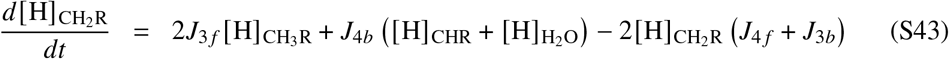

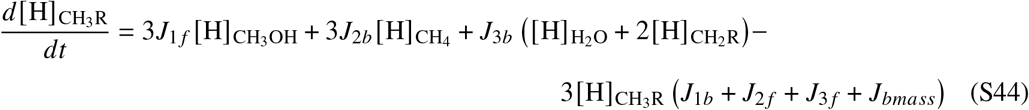

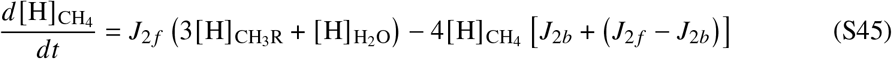

We isolate the terms for 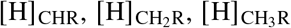, and 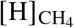 under a steady state assumption. Given *r*_*i*_ = *J*_*ib*_/*J*_*i f*_ we define the following identities:

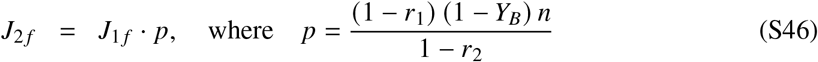

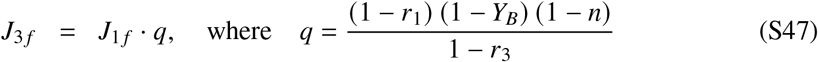

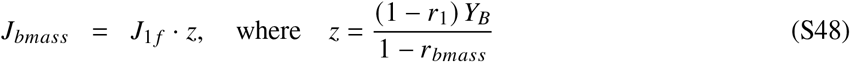

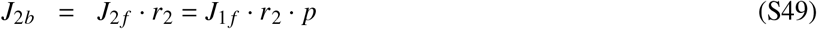

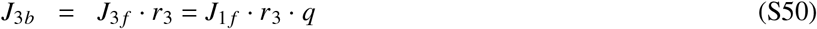

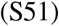

We now isolate the terms for 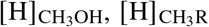, and 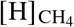, and define *j* and *k*:

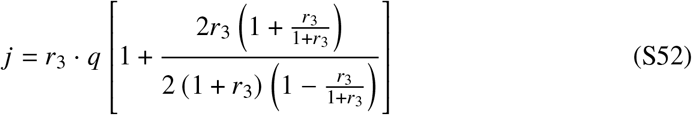

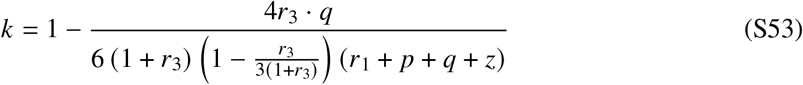

Expressing Eq. S45 in terms of *p, q, z, j*, and *k* yields:

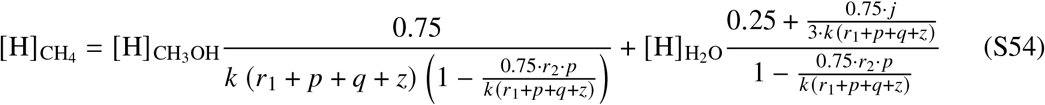

We define the right term in the RHS as *x*_*w*_, the fraction of the water-derived hydrogen atoms in CH_4_, and separate *x*_*w*_:

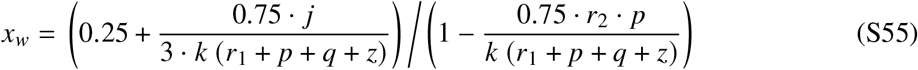

We followed a similar approach for the acetoclastic pathway. Here *n* = 1, and the cycling term *m* is defined similarly to the isotopic model. Following the derivation for the isotopes where we assumed that CH_2_R is at equilibrium with water, any hydrogen that goes back from CH_2_R to CH_3_R originates from water.

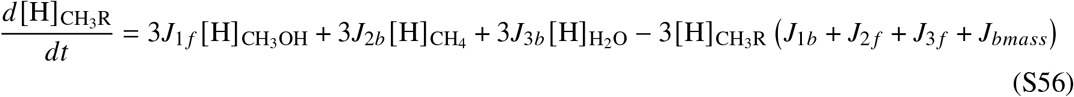

and rearrange to isolate *x*_*w*_:

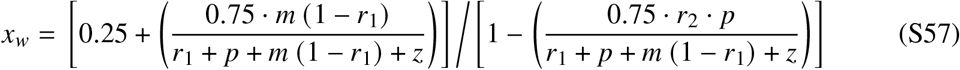

### Supplementary Text

#### Fitted Hydrogen Kinetic Isotopic Effects

We discuss here the median values of the posterior hydrogen KIE distributions, and define the uncertainties as 95% of the model results (Table S6). As a consequence of the reaction network structure, the model is the most sensitive to the KIEs of CH_3_-H_4_MPT:coenzyme M methyltransferase (MTR) and MCR, which yields a relatively narrow uncertainty in the posterior distributions. The secondary and primary KIEs of MCR here were 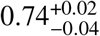and 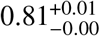, respectively. The only measurement of MCR hydrogen KIEs are from a thermophilic methanogenic strain (*Methanother - mobacter marburgensis*) grown at 60 °C, with a secondary KIE of 0.84 ± 0.01 and a primary KIE of 0.77 ± 0.2. While we culture a different, mesophilic methanogen, our fitted KIEs are well within the large uncertainty range of the measured primary KIE of MCR. Our fitted secondary KIE is slightly larger than the measured secondary KIE, which agrees with the general prediction that KIE increase in lower temperatures (*32*). The secondary KIE of MTR in the direction of CH_3_-S-CoM oxidation is 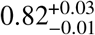. This KIE was not measured before, but was predicted to be 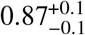 in a model of hydrogenotrophic methanogenesis (*12*), and our predicted value is within the uncertainty range. The remaining KIEs in the model have larger uncertainties, due to their smaller contribution in hydrogen isotopic exchange in the baseline scenario on WT growth. The primary KIEs of MER, MTD, and formyl methanofuran dehydrogenase (FMD) are 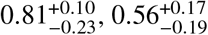, and 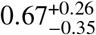, respectively, and the secondary KIEs of MER and MTD are 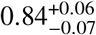and 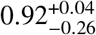, respectively. The large uncertainties for these reactions reflect the model’s insensitivity for their values in the WT growth scenario.

#### Carbon Isotopic Composition of Methane

In the main text we interpreted the *δ*^13^C-CH_4_ values at the end of the growth experiments in the context of the isotopic model. We observed that during growth on methanol, *δ*^13^C-CH_4_ values in the lowest *mcr* expression levels (i.e., no tetracycline added) were significantly lower than the wild-type by 5 ‰ (*p* < 0.01, figure S2). Following the hypothesis that this change is related to increased reversibility of the oxidative branch or MCR, using the known equilibrium isotopic effects (EIEs) for these reactions could inform us on the associated predicted directions of change in *δ*^13^C-CH_4_. The oxidative branch has a large and negative EIE between CH_3_-S-CoM and CO_2_ (−64 ‰ at 37 ^°^C), while MCR has a small positive EIE between CH_3_-S-CoM and CH_4_ (1 ‰ at 37 ^°^C). In other words, under isotope equilibrium, the CO_2_ is expected to be isotopically heavier than the CH_3_-S-CoM, which CH_4_ would be slightly isotopically lighter. MCR has a normal (i.e, *α* < 1) carbon kinetic isotopic effect (KIE) in the direction of methane production [≈−40 ‰, Ref. (*18*)]. Based on a different isotopic-enabled metabolic model, MTR, the first reaction in the oxidative branch has a predicted normal carbon KIE [≈−11 ‰, Ref. (*12*)]. We note that both these values are from hydrogenotrophic methanogens and are only used here for illustrative purposes. In wild-type growth, where we predict there is only minimal isotopic exchange (Fig. 1B), the relation of these two KIEs determine the isotopic fractionation in the branching point where CH_3_-S-CoM is partially reduced to methane, and partially oxidized to CO_2_ (*33*). At the end of an experiment, we expect that the total *δ*^13^C-CH_4_ and *δ*^13^C-CO_2_ weighted by molar quantities would be −41.6 ‰ (based on the initial *δ*^13^C of the methanol). The *δ*^13^C-CH_4_ values in the wild-type are ≈8 ‰ more negative (*δ*^13^C-CH_4_ values of −49.2 ‰), consistent with previous experiments (*34, 35*). Following the mass balance *δ*^13^C-CH_3_OH = 0.75 · *δ*^13^C-CH_4_ + 0.25 · *δ*^13^C-CO_2_ (and ignoring the biomass), we can calculate the expected *δ*^13^C-CO_2_ to be −18.8 ‰. The isotopic fractionation between CH_4_ and CO_2_ determined by *α* = (1000 + *δ*^13^C-CH_4_)/(1000 + *δ*^13^C-CO_2_) is 0.969. At a steady-state (i.e., no change of the isotopic compositions and concentrations with time), we can write:

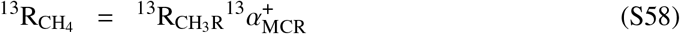

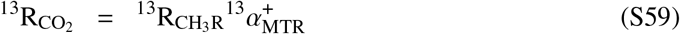

where ^13^R_i_ is the ^13^C/^12^C ratio of compound *i*, and 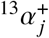is the forward KIE of reaction *j*. We can now express the isotopic fractionation between CH_4_ and CO_2_ in terms of the KIEs:

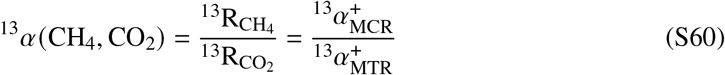

Taking the aforementioned KIEs for MCR and MTR from hydrogenotrophic methanogenesis we obtain 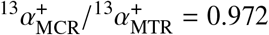, matching well our predicted ^13^*α*(CH_4_, CO_2_) of 0.969 in the WT methylotrophic growth conditions.

Now we turn to the predicted effects in the case of isotopic exchange in MCR or the oxidative branch. An increasing degree of reversibility in those pathways will shift the observed net isotopic effect toward the EIEs (*36*). In the first scenario, increased reversibility of only MCR would shift the apparent KIE of the reaction from ≈−40 ‰ to ≈1 ‰, decreasing the difference between the isotopic effects in the oxidative branch and leading to a more positive *δ*^13^C-CH_4_. In the second scenario, increased reversibility of only the oxidative branch would shift the apparent KIE of the reaction from ≈−11 ‰ to ≈−64 ‰, further increasing the difference between the isotopic effects in the oxidative branch, leading to a more negative *δ*^13^C-CH_4_. In other words, expressing the large and negative EIE of the oxidative branch causes a decrease *δ*^13^C of CH_3_-S-CoM, which translates to a more negative *δ*^13^C-CH_4_ value, which we observed. We conclude that this simplified analysis predicts that increased carbon isotopic exchange in the oxidative branch would lead to more negative *δ*^13^C-CH_4_ values.

During growth on acetate, the *δ*^13^C-CH_4_ values at the end of the experiment (from −38.4 ‰ to −37.2 ‰) were similar to the *δ*^13^C value of the methyl group of the acetate (−37.1 ‰). This is in agreement with the acetoclastic metabolic pathway were the methyl group is fully converted to methane (minus the biomass). The small differences that we did observe in our experiments (≈1 ‰) might be attributed to isotopic exchange in the cycling reaction, which we included in our hydrogen isotopic model. This reaction does not generate CO_2_, but could allow some carbon isotopic exchange between CH_3_-S-CoM and other more oxidized intermediates that are required for anabolism. We conclude that our experimental results, which show a only a minor change in *δ*^13^C-CH_4_ with decreasing *mcr* expression levels, match our current understanding of the acetoclastic pathway.

**Figure S1:**
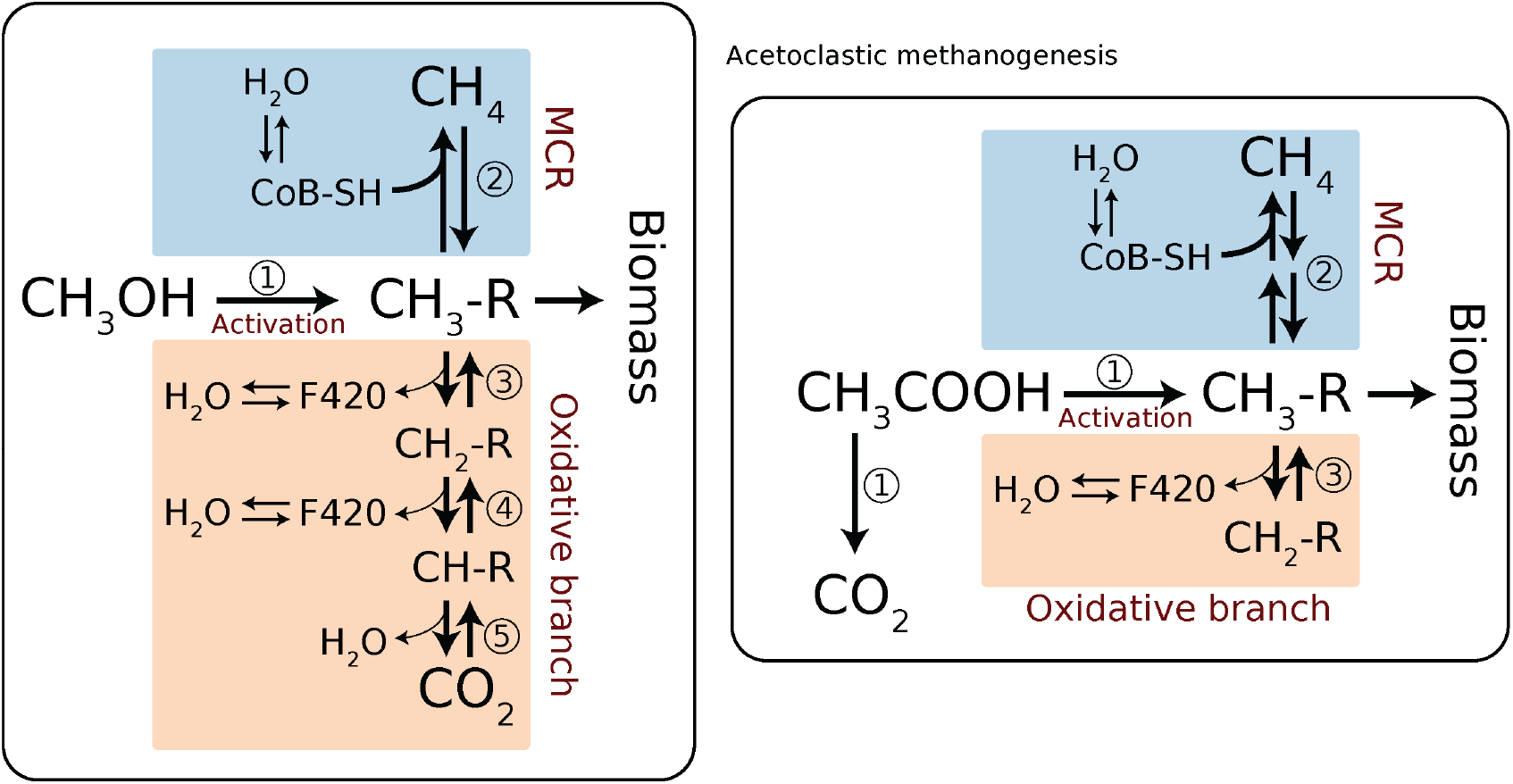
Reactions included in the isotopic mass balance for methylotrophic methanogenesis with methanol (left) and acetoclastic methanogenesis (right). For simplicity, we only include the oxidation/reduction reactions, and abbreviate the full metabolite names to highlight the redox status of the carbon. The MCR catalyzed reaction (#2) exchanges hydrogen with HS-CoB, the MER and MTD catalyzed reactions (#3 and #4) exchange hydrogen with coenzyme F420, and the FMD catalyzed reaction exchanges hydrogen directly with H_2_O. We assume here that the H_2_O pool is infinite so that its *δ*Dis constant during the exeriment. We also assume that HS-CoB and F420 are at isotopic equilibrium with the H_2_O (see text for further explanation).

**Figure S2:**
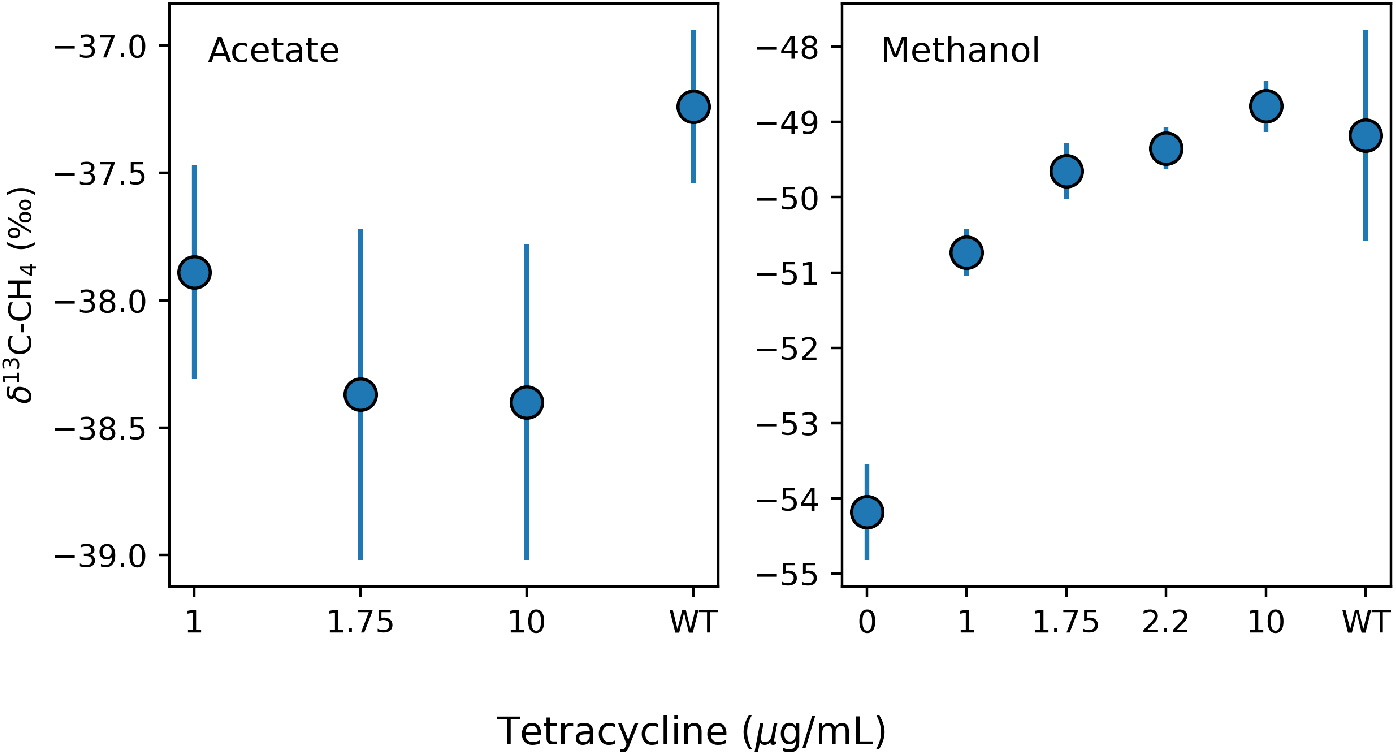
The carbon isotopic composition of methane (*δ*^13^C-CH_4_) at the end of the growth experiments on acetate (left) and methanol (right), as a function of tetracycline concentration. The error bars represent the standard deviation (1*σ*) for 4 replicates grown on medium that contains different levels of deuterated H2O (*δ*D-H_2_O between *approx* −90 ‰ and ≈ +2200 ‰).

**Figure S3:**
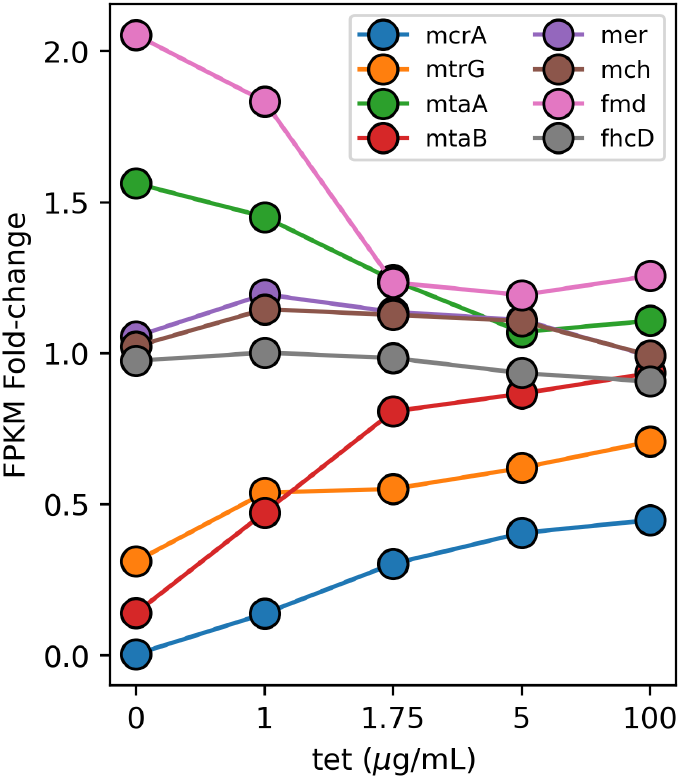
Fold change of RNA transcripts for a DDN032 on various tetracycline induction levels relative to the wild-type strain of *M. acetivorans* grown on methanol. Data from Ref. (*2*).

**Figure S4:**
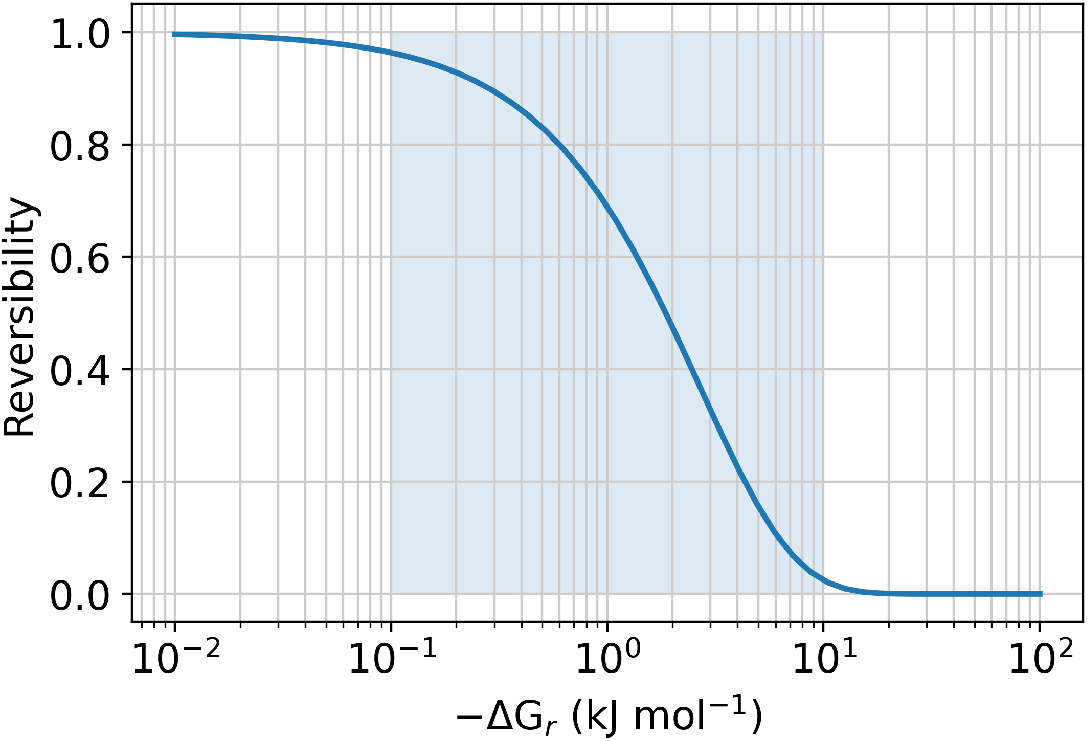
The reversibility of a chemical reaction as a function of its Gibbs free energy (ΔG_*r,i*_). The reversibility is defined as the ratio of the backward to forward gross fluxes (*r*_*i*_ = *J*_*i,b*_/*J*_*i, f*_), and is related to the ΔG_*r,i*_ via *r*_*i*_ = exp (ΔG_*r,i*_/RT) ranging between 0 (irreversible) to 1 (fully reversible). Due to the exponential relation between the reversibility and the ΔG_*r,i*_, a very narrow range of ΔG_*r,i*_ between ≈−10 kJ mol^−1^ and ≈−0.1 kJ mol^−1^ (represented by the highlighted area) corresponds to drastic changes in the reversibility from ≈5% to ≈95%. A key takeaway is that at ΔG_*r,i*_ values more negative than −10 kJ mol^−1^ would practically have a only minor effect on the reversibility.

**Figure S5:**
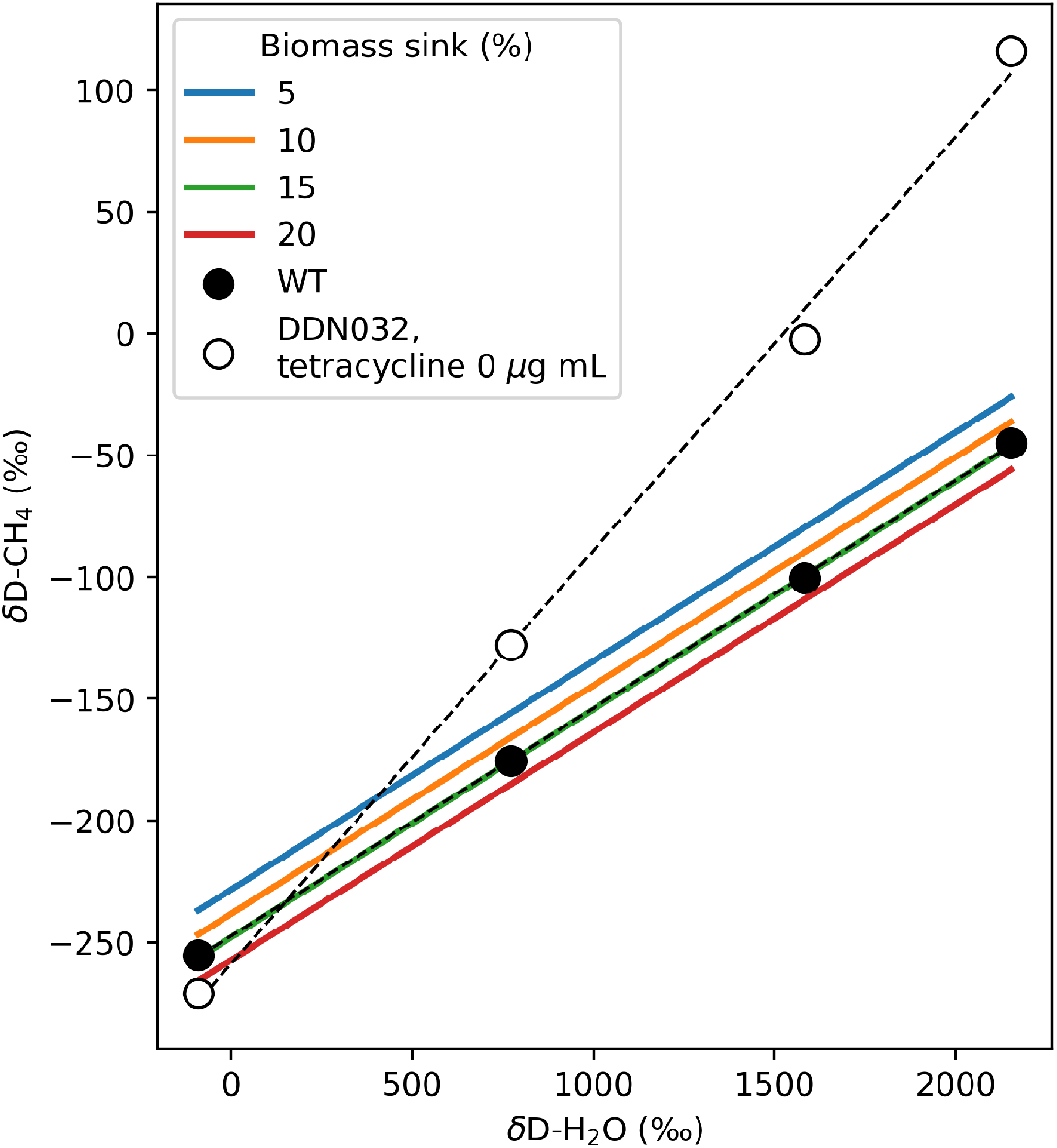
Sensitivity of the isotopic model of methylotrophic methanogenesis to the magnitude of the biomass sink. In the model we used a biomass sink value of 15%. Changes in the biomass sink magnitude induce changes in the y-intercept of *δ*D-CH_4_ vs. *δ*D-H_2_O. The measured *δ*D-CH_4_ values of the wild-type *M. acetivorans* and the DDN032 mutant grown on 0 *μ*g/mL tetracycline are shown for reference.

**Table S1:** Growth rates of mutant and wild-type *M. acetivorans* strains under different tetracycline induction levels. Growth rates for growth on methanol were not monitored and were estimated based on Ref. (*2*).

| Strain | Substrate | Tetracycline<br>($\mu\text{g/mL}$) | Growth rate<br>($\text{h}^{-1}$) | $\pm (1\sigma)$ | Doubling time (h) | $\pm (1\sigma)$ | $n$ |
| --- | --- | --- | --- | --- | --- | --- | --- |
| DDN032 | Acetate | 0 | 0.0018 | 0.0007 | 440 | 183 | 3 |
| DDN032 |  | 1 | 0.0024 | 0.0006 | 305 | 70 | 3 |
| DDN032 |  | 1.75 | 0.0057 | 0.0016 | 129 | 31 | 4 |
| DDN032 |  | 10 | 0.0105 | 0.0025 | 69 | 16 | 4 |
| WWM60 |  | – | 0.0107 | 0.0011 | 65 | 6 | 3 |

**Table S2:** Measured isotopic compositions of methane (*δ*D-CH_4_ and *δ*^13^C-CH_4_) and water (*δ*D-H_2_O) for *M. acetivorans* growth on methanol. The water and CH_4_ were sampled at the end of the growth experiments. *δ*D values are relative to VSMOW, and *δ*^13^C values are relative to VPDB.

| Sample | Tetracycline<br>($\mu\text{g/mL}$) | $\delta\text{D-H}_2\text{O}$ (‰) | $\delta\text{D-CH}_4$ (‰) | $\delta^{13}\text{C-CH}_4$ (‰) |
| --- | --- | --- | --- | --- |
| 0 | 0 | -90.1 | -270.8 | -54.69 |
|  | 0 | 772.4 | -128.0 | -54.50 |
|  | 0 | 1582.0 | -2.4 | -53.25 |
|  | 0 | 2153.3 | 115.9 | -54.28 |
| 1 | 1 | -90.1 | -267.2 | -51.16 |
|  | 1 | 772.4 | -141.6 | -50.74 |
|  | 1 | 1582.0 | -34.3 | -50.55 |
|  | 1 | 2153.3 | 39.2 | -50.47 |
| 1.75 | 1.75 | -90.1 | -263.128 | -49.36 |
|  | 1.75 | 772.4 | -156.7 | -49.33 |
|  | 1.75 | 1582.0 | -51.1 | -49.86 |
|  | 1.75 | 2153.3 | 19.2 | -50.07 |
| 2.2 | 2.2 | -90.1 | -256.1 | -43.58<sup>†</sup> |
|  | 2.2 | 772.4 | -158.4 | -49.55 |
|  | 2.2 | 1582.0 | -55.9 | -42.04<sup>†</sup> |
|  | 2.2 | 2153.3 | 14.8 | -49.15 |
| 10 | 10 | -90.1 | -260.3 | -48.60 |
|  | 10 | 772.4 | -166.7 | -49.30 |
|  | 10 | 1582.0 | -66.2 | -48.70 |
|  | 10 | 2153.3 | -21.8 | -48.57 |
| WT | 0 | -90.1 | -255.3 | -50.26 |
|  | 0 | 772.4 | -175.4 | -48.09 |
|  | 0 | 1582.0 | -100.5 | -50.51 |
|  | 0 | 2153.3 | -44.9 | -47.84 |

<sup>†</sup> Samples that showed anomalously high $\delta^{13}\text{C-CH}_4$ values. We suspect that those samples contained small amounts of $\text{N}_2$ or $\text{O}_2$ which interfered with the measurements. We this did not include these data points in figure S2.

**Table S3:** Measured isotopic compositions of methane (*δ*D-CH_4_ and *δ*^13^C-CH_4_) and water (*δ*D-H_2_O) for *M. acetivorans* growth on acetate. The water and CH_4_ were sampled at the end of the growth experiments. *δ*D values are relative to VSMOW, and *δ*^13^C values are relative to VPDB.

| Sample | Tetracycline<br>($\mu\text{g/mL}$) | $\delta\text{D-H}_2\text{O}$ (‰) | $\delta\text{D-CH}_4$ (‰) | $\delta^{13}\text{C-CH}_4$ (‰) |
| --- | --- | --- | --- | --- |
| 1 | 1 | -91.6 | -340.4 | -38.10 |
|  | 1 | 718.8 | -177.1 | -37.82 |
|  | 1 | 1503.6 | -6.2 | -38.31 |
|  | 1 | 1925.2 | 41.4 | -37.33 |
| 1.75 | 1.75 | -91.6 | -336.90 | -39.26 |
|  | 1.75 | 718.8 | -196.7 | -37.71 |
|  | 1.75 | 1503.6 | -92.5 | -38.24 |
|  | 1.75 | 1925.2 | 12.2 | -38.28 |
| 10 | 10 | -91.6 | -332.4 | -39.29 |
|  | 10 | 718.8 | -218.6 | -37.84 |
|  | 10 | 1503.6 | -116.2 | -38.14 |
|  | 10 | 1925.2 | -70.1 | -38.34 |
| WT | 0 | -91.6 | -324.1 | -37.55 |
|  | 0 | 815.1 | -219.8 | -37.33 |
|  | 0 | 1428.1 | -156.9 | -37.26 |
|  | 0 | 1947.0 | -108.7 | -36.82 |

**Table S4:** Linear regression parameters between *δ*D-CH_4_ and *δ*D-H_2_O for growth on methanol and acetate in different *mcr* induction levels. The slope and regression are calculated by the *δ*D/1000+1 values.

| Strain | Substrate | Tetracycline<br>($\mu\text{g/mL}$) | Slope | ± (1SE) | Intercept<br>($\delta\text{D}/1000+1$) | ± (1SE) | *n* |
| --- | --- | --- | --- | --- | --- | --- | --- |
| DDN032 | Methanol | 0 | 0.170 | 0.007 | 0.498 | 0.017 | 4 |
| DDN032 |  | 1 | 0.136 | 0.003 | 0.595 | 0.014 | 4 |
| DDN032 |  | 1.75 | 0.126 | 0.001 | 0.605 | 0.016 | 4 |
| DDN032 |  | 2.2 | 0.121 | 0.002 | 0.616 | 0.017 | 4 |
| DDN032 |  | 10 | 0.109 | 0.005 | 0.629 | 0.023 | 4 |
| WWM60 |  | – | 0.094 | 0.001 | 0.645 | 0.011 | 7 |
| DDN032 | Acetate | 1 | 0.195 | 0.019 | 0.492 | 0.002 | 3 |
| DDN032 |  | 1.75 | 0.167 | 0.013 | 0.511 | 0.006 | 4 |
| DDN032 |  | 10 | 0.131 | 0.007 | 0.553 | 0.003 | 3 |
| WWM60 |  | – | 0.106 | 0.004 | 0.585 | 0.015 | 7 |

**Table S5:** Standard transformed Gibbs free energies of the methylotrophic and acetoclastic pathways. Calculated by eQuilibrator 3.0 (*37*).

| | Reaction | $\Delta G_r'^0$<br>(kJ mol<sup>-1</sup> CH<sub>4</sub>) |
| --- | --- | --- |
| **Net methylotrophic** | $\text{CH}_3\text{OH} \rightleftharpoons 0.75\text{CH}_4(\text{aq}) + 0.25\text{CO}_2(\text{aq}) + 0.5\text{H}_2\text{O}(\text{l})$ | $-88.1 \pm 9.9$ |
| MTA | $\text{CH}_3\text{OH} + \text{HS-CoM} \rightleftharpoons \text{H}_2\text{O}(\text{l}) + \text{CH}_3\text{-SCoM}$ | $-57.3 \pm 9.9$ |
| MCR | $0.75\text{CH}_3\text{SCoM} + 0.75\text{HS-CoB} \rightleftharpoons$<br>$0.75\text{CH}_4(\text{aq}) + 0.75\text{CoM-SS-CoB}$ | $-24.9 \pm 8.1$ |
| Oxidative branch | $0.25\text{CH}_3\text{-SCoM} + 0.75\text{CoM-SS-CoB} + 0.5\text{H}_2\text{O}(\text{l}) \rightleftharpoons$<br>$0.75\text{HS-CoB} + \text{HS-CoM} + 0.25\text{CO}_2(\text{aq})$ | $-5.9 \pm 4.8$ |
| **Net acetoclastic** | $\text{CH}_3\text{COO}^- \rightleftharpoons \text{CH}_4(\text{aq}) + \text{CO}_2(\text{aq})$ | $-7.6 \pm 8.4$ |
| ACK/PTA/ACDS | $\text{CH}_3\text{COO}^- + \text{H}_4\text{MPT} + \text{CoM-SS-CoB} \rightleftharpoons$<br>$\text{CH}_3\text{-H}_4\text{MPT} + \text{HS-CoM} + \text{HS-CoB} + \text{CO}_2(\text{aq})$ | $+59.0 \pm 13.2$ |
| MTR/MCR | $\text{CH}_3\text{-H}_4\text{MPT} + \text{HS-CoM} + \text{HS-CoB} \rightleftharpoons$<br>$\text{CH}_4 + \text{CoM-SS-CoB} + \text{H}_4\text{MPT}$ | $-66.6 \pm 13.0$ |

**Table S6:** The posterior hydrogen isotope kinetic isotope effects (KIEs) fitted with data from growth of the wild-type strain on methanol and acetate. The *s* and *p* subscripts denote secondary and primary kinetic isotope effects, respectively. The range of uncertainties represents 95% of the model results. Abbreviations: H_4_MPT, tetrahydromethanopterin; MTR, methyl-H_4_MPT: coenzyme M methyltransferase; MCR, methyl coenzyme M reductase; MER, methylene H_4_MPT reductase; MTD, F420-dependent methylene H_4_MPT dehydrogenase; FMD, formyl methanofuran dehydrogenase.

| Enzyme | KIE<sub>s</sub> | KIE<sub>p</sub> |
| --- | --- | --- |
| MTR | 0.82<sup>+0.03</sup><sub>-0.01</sub> | — |
| MCR | 0.74<sup>+0.02</sup><sub>-0.04</sub> | 0.81<sup>+0.01</sup><sub>-0.00</sub> |
| MER | 0.84<sup>+0.06</sup><sub>-0.07</sub> | 0.81<sup>+0.10</sup><sub>-0.23</sub> |
| MTD | 0.92<sup>+0.04</sup><sub>-0.26</sub> | 0.56<sup>+0.17</sup><sub>-0.19</sub> |
| FMD | — | 0.67<sup>+0.26</sup><sub>-0.35</sub> |

